# Deterministic and stochastic rules of branching govern dendritic morphogenesis of sensory neurons

**DOI:** 10.1101/2020.07.11.198309

**Authors:** Amrutha Palavalli, Nicolás Tizón-Escamilla, Jean-François Rupprecht, Thomas Lecuit

**Affiliations:** Aix Marseille Université & CNRS, IBDM - UMR7288 & Turing Centre for Living Systems Campus de Luminy Case 907, 13288, Marseille, France; Aix-Marseille Université, Université de Toulon, CNRS, CPT, Turing Centre for Living Systems, Marseille, France; Collège de France, 11 Place Marcelin Berthelot, 75005 Paris

## Abstract

Dendrite morphology is necessary for the correct integration of inputs that neurons receive. The branching mechanisms allowing neurons to acquire their type-specific morphology remain unclear. Classically, axon and dendrite patterns were shown to be guided by molecules providing deterministic cues. However, the extent to which deterministic and stochastic mechanisms, based upon purely statistical bias, contribute to the emergence of dendrite shape is largely unknown. We address this issue using the *Drosophila* class I vpda multi-dendritic neurons. Detailed quantitative analysis of vpda dendrite morphogenesis indicates that the primary branch grows very robustly in a fixed direction while secondary branch numbers and lengths showed fluctuations characteristic of stochastic systems. Live tracking dendrites and computational modeling revealed how neuron shape emerges from few local statistical parameters of branch dynamics. We report key opposing aspects of how tree architecture feedbacks on the local probability of branch shrinkage. Child branches promote stabilization of parent branches while self-repulsion promotes shrinkage. Finally, we show that self-repulsion, mediated by the adhesion molecule Dscam1, indirectly patterns the growth of secondary branches by spatially restricting their direction of stable growth perpendicular to the primary branch. Thus, the stochastic nature of secondary branch dynamics and the existence of geometric feedback emphasizes the importance of self-organization in neuronal dendrite morphogenesis.

## Introduction

Dendrites are neuronal processes that are specialized in receiving information. They have strikingly different arborization patterns depending on neuronal subtypes. Dendrite arborization patterns depend on several factors such as the number and types of synaptic or sensory inputs received by the neuron or the geometry and size of the receptive fields. Failure to establish proper branching patterns and circuitry leads to various neurodevelopmental and neuropsychiatric disorders ^1^.

In general, dendrite morphogenesis follows a set of consecutive steps, which include (i) dendrite initiation, (ii) outgrowth (iii) branching and maturation (iv) establishment of boundaries and in a few cases, arbor remodeling ^2,3^. Each of these processes requires extensive regulation by cell intrinsic cues as well as signals coming from its environment.

Dendrite morphology is regulated cell-intrinsically by transcription factors ^4–8^, cytoskeletal regulators ^9–15^, components of endocytic pathway ^16–18^ and secretory pathway ^19,20^ all of which have been shown to either promote or reduce dendrite branching.

Extrinsic cues include, long-range diffusible secreted factors that are classically involved in axon guidance like Semaphorins ^21^, Slits ^22–24^ and Netrins ^25–27^ have also been shown promote or restrict dendrite branching. Contact mediated, short range cues like cell adhesion molecules, adhesion GPCRs guide branch points ^28^, restrict branching ^29^ and maintain dendrites in a 2D plane ^30^. Neuronal activity also refines arbor morphology by increasing or decreasing branching density ^31,32^. Though much is known about the molecules regulating dendrite branching, how they govern the neuron-specific arborization pattern remains unclear. Indeed, most molecules tend to increase or decrease branching density without significantly affecting arborization patterns.

Two modes of branching morphogenesis have emerged from studying other branched organ systems ^33–35^. The first mode is a deterministic one, where systems have highly stereotyped branching patterns as described in the mouse lung ^36^ and *Drosophila* tracheal system ^37^. In deterministic systems, branching is orchestrated by patterned cues such as *Drosophila* FGF, which is expressed in clusters of cells surrounding the tracheal sacs at specific positions and instructs the branch points of the primary bud^37^. The dendrites of the *C. elegans* PVD neuron represent an extreme example of deterministic patterning in the neuronal system. The branch points in the complex menorah-like PVD dendrites are almost exclusively determined by patterned cues in the epidermis^28,38–40^. The second mode of branching is self-organized, wherein final morphology emerges from the statistical features of branch dynamics and the local interactions between growing tips. This results in stochastic branch patterns as described in the mammary gland^34^. Although extrinsically derived FGF promotes branching in this system, no patterned cue has been identified that could potentially guide branching^41,42^. The branching features of the mammary gland can be recapitulated by a model that accounts for local parameters such as tip elongation rate, branching rate and mode of tip termination^34^. Thus, branching morphogenesis of the mammary gland follows a stochastic, self-organized scheme and does not seem to require extrinsically patterned cues.

In the neuronal system, cell fate has traditionally been thought to be achieved deterministically by specific regulatory genes^43,44^ but neuronal connectivity is refined by an activity-dependent, self-organization^45^. However, the relative contribution of deterministic and self-organized mechanisms in establishing dendrite patterns remains unknown.

The *Drosophila melanogaster* multi dendritic-dendritic arborization (md-da) neurons are the model of choice to address this question. The md-da neurons are part of the peripheral nervous system and are involved in somatosensation. They are divided into four distinct morphological classes in an increasing order of dendritic complexity. They exhibit stereotyped dendritic structures that are identifiable across animals and are restricted in a 2-D space beneath the epidermis ^46–49^. Md-da neurons have been extensively used to study dendrite morphogenesis, especially class I and class IV neurons which exhibit the simplest and most complex morphologies respectively ^6,8,50,51^. However, most of these studies were based on fixed imaging and have focused on late developmental stages, after the establishment of the typical dendritic morphology of the four neuronal classes. Two studies have described the embryonic development of the dorsal class I neurons ^52,53^ but they were mainly qualitative and lacked appropriate temporal resolution.

In this study, we use high-resolution live imaging to quantitatively describe morphogenesis of class I vpda neuron. We show that the primary dendrites grow deterministically while secondary dendrite patterning is stochastic. Additionally, our computational model and Dscam1 data shows that self-repulsion patterns secondary dendrites by preferentially stabilizing them orthogonally from the primary dendrite giving the class I neurons their characteristic morphology.

## Results

### Class I vpda neuron shape is established during embryogenesis

In order to understand how class I specific dendrite morphology is achieved; we first investigated when the final morphology of their dendritic arbor is established. The vpda neuron has one large primary dendrite that projects dorsally and another small primary branch on the ventral side. Secondary dendrite project outwards from the side of the primary dendrites giving the class I neurons their characteristic “bottle brush” morphology. Our study focused on the dorsal dendrite of the vpda neuron since its location was convenient for whole mount imaging.

Dendrite morphology was analyzed at 4 developmental time points i.e. late embryogenesis, (E17, 19±2 hours After Egg Laying, or h AEL), 1st instar (L1, 24±3h AEL), 2nd instar (L2, 48±3h AEL) and 3rd instar (L3, 72±3h AEL) (Fig. 1a). Neurons were labeled with UAS-mCD8::GFP, expressed by the neuronal class I specific driver 2-21gal4 ^5^. Consistent with the previous descriptions of the dorsal class I neurons ^52^, we observed a qualitative enlargement of ventral vpda neurons over time that correlated with the growth of the organism (Fig. 1a). To confirm this, we developed tools to quantitatively describe dendrite morphology. Dendrites of vpda neurons were segmented and classified into primary, secondary, tertiary and quaternary branches (Schematic Fig. 1a and Fig. S1a-c). The branch numbers for each order of complexity were first measured (Fig. 1b). On average, 15-17 secondary branches emanated from the primary branch across the four developmental time points showing no statistically significant changes occurred during development (Fig. 1b). The primary and secondary branches made up the bulk of the dendrite length (84.6% on average) at L3 (Fig. S1d). The tertiary and quaternary branches made up, on average, 15.4% of the total dendrite length at L3 (Fig. S1d), and showed some changes during development but did not significantly alter the global dendrite pattern.

**Figure 1.**
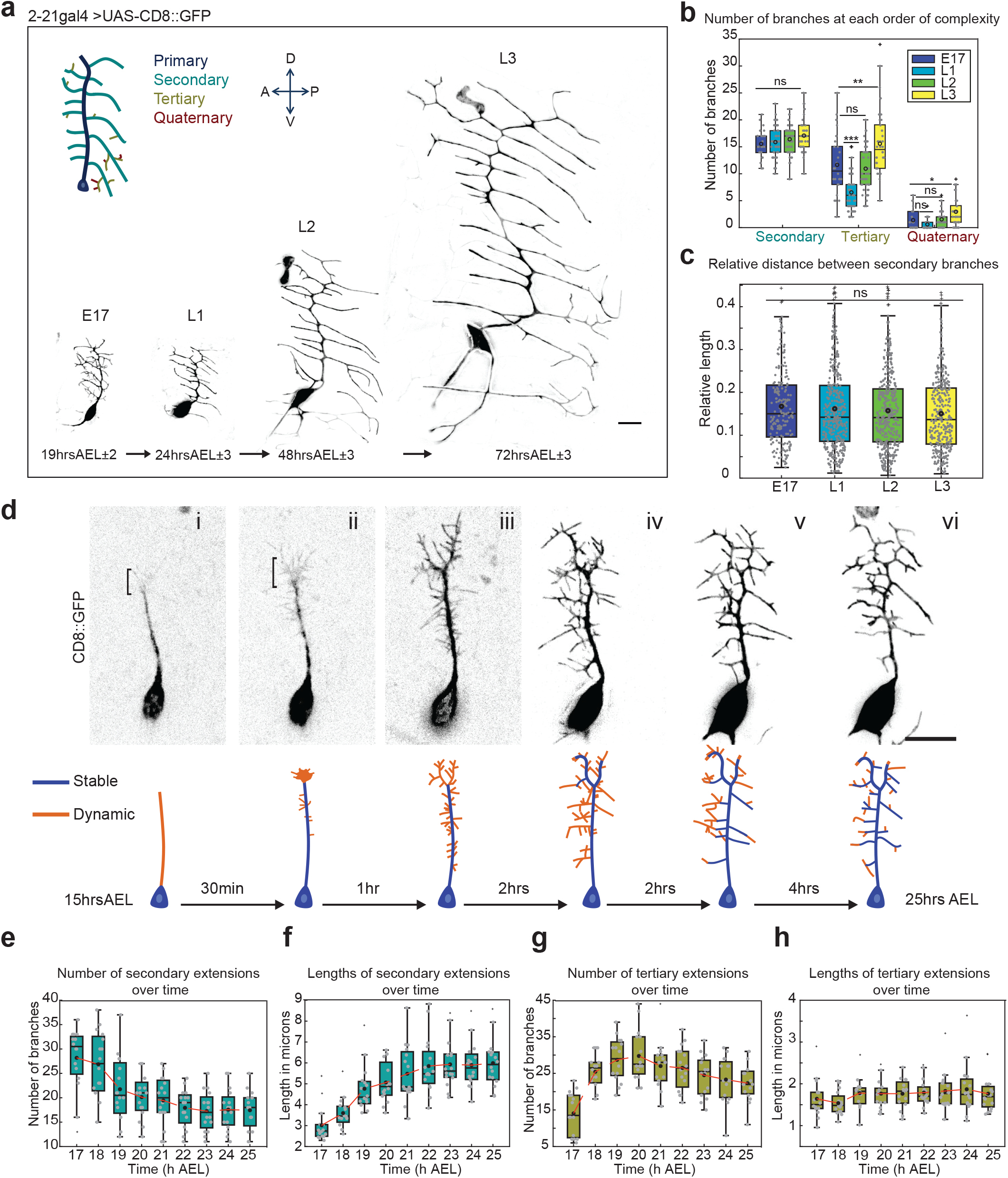
Dendrite patterning is nearly complete by the end of embryogenesis. (a) Images of Class I vpda neurons taken from homozygous 2-21gal4/UAS-mCD8::GFP embryos and larvae at the following stages: late embryo (E17 19h AEL±2), 1st instar (L1 24h AEL±3), 2^nd^ instar (L2 48h AEL±3), 3^rd^ instar (L3 72h AEL±3) Schematic: Primary branch (blue) Secondary branches (cyan) Tertiary (brown) Quaternary (red). (b) Quantification of the number of branches at each order of complexity in the neuron. E17 n=26, L1 n=48, L2 n=50 L3 n=38 neurons. (c) Relative distance (Inter dendrite distance/primary branch length) across the 4 stages. E17 n=194, L1 n=374, L2 n=409 L3 n=341 inter-dendrite segments (d) Top panel: Time-lapse images of dendrite development of 2-21gal4/UAS-mCD8::GFP mhc1 mutant background embryos from ~15h AEL to ~25h AEL (Movie1). Lower panel: Schematic representation of dendrite development. Indigo represents stable dendrites and orange represents dynamic extensions. (e) Number of Secondary Branches from 17-25h AEL. (f) Lengths of Secondary Branches from ~17-25h. (g) Number of Tertiary Branches from 17-25h AEL. (h) Length of Tertiary Branches from ~17-25h AEL for (e)-(h) AEL n=16 neurons. For each box, the central line is the median, the ‘o’ is the mean, the box extends vertically between the 25th and 75th percentiles, the whiskers extend to the most extreme data that are not considered outliers, and the outliers are plotted individually. Statistical significance has been calculated using Mann-Whitney U test for (b) and (c) Not significant= ns, * p<0.05; ** p<0.01, *** p<0.001. All the panels have the same orientation: dorsal at the top, anterior to the left. Scale bars = 10μm.

The total dendrite lengths made up by the primary, secondary, tertiary and quaternary branches also did not vary greatly across all 4 developmental stages (Fig. S1d). Thus, the core morphology of vpda neurons dendrites is established in the embryo. Consistent with this, the relative distance between secondary branches did not change over time (Fig. 1c), thus confirming that the dendritic pattern scales isometrically in size as the larva grows.

To assess branching mechanisms controlling neuron morphology, we set out to image dendrite morphogenesis in living embryos. Using the setup described in the methods section we obtained movies covering the entire dendrite morphogenesis (Movie1). These movies reveal 3 phases in vpda neuron morphogenesis: *(i) Primary dendrite formation*: Between 13-15h AEL, vpda neurons extend a single and stable primary dendrite towards the dorsal region of the embryo (Fig. 1d, Movie2), similar to the dorsal class I neurons, described previously ^52,53^. *(ii) Secondary branch initiation and elongation:* After ~15h AEL, flat lamellipodia-like membrane protrusions started appearing at the distal tip of the primary dendrite (Fig. 1d (i), (ii)). At ~17h AEL, numerous, small and very dynamic protrusions that constantly extend and retract appeared everywhere along the primary dendrite (Fig. 1d (iii)). These dynamic protrusions were termed secondary extensions to distinguish them from the comparatively stable secondary branches observed at later stages. On average 28.2±6.4 secondary extensions were present at 17h AEL (Fig. 1e). Over the next 3-4 hours these extensions elongated at a rate of ~1μm/hour (Fig. 1f). The growth of secondary extensions was accompanied by the emergence of dynamic protrusions branching off from secondary extensions, termed tertiary extensions. *(iii) Secondary branch stabilization:* After 20h AEL the dendritic structure became markedly less dynamic and secondary extension lengths (6±1.2μm) and numbers (17.6±3) stabilized at 22h AEL (Fig. 1 e-f). On average 62.3% of the initial secondary extensions were stabilized as secondary dendrites, suggesting that the secondary extensions may be in competition to form stable dendrites (Fig. 1e). The number of tertiary extensions dropped as the secondary extensions stabilized into secondary dendrites (Fig. 1f-g). This number is even lower at later L1 (Fig. 1b) suggesting that the tertiary extensions are transient structures. Unlike secondary extensions no drastic change in tertiary extension lengths was observed over time (Fig. 1h).

In summary, the primary branch grows dorsally in a straightforward manner followed by the emergence of dynamic secondary extensions undergoing repeated cycles of extension and retraction until stabilization of a subset of branches. Our observations therefore reveal striking differences in the mode of development of the primary and secondary dendrites.

### Growth of the primary dendrites is deterministic

Every class I vpda neuron consistently forms a single dorsal primary dendrite that extends towards the dorsal region of the embryo (Fig. 2a, Movie2). This reproducible growth pattern suggested a stringent control of primary dendrite number and growth direction and hence deterministic growth.

**Figure 2.**
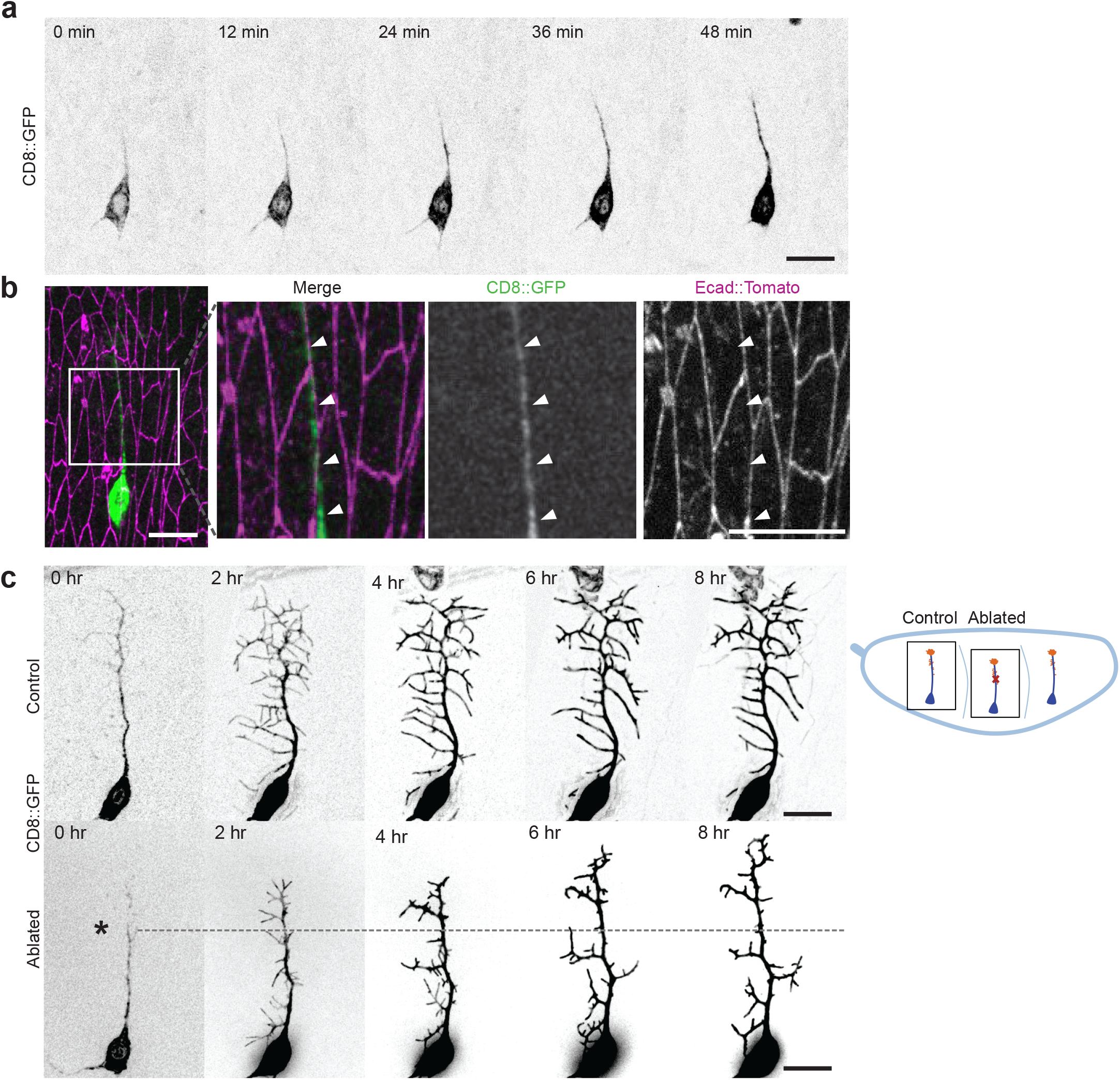
Development of the primary branch is consistent with deterministic growth. (a) Time-lapse images of the uni-directional growth of the primary dendrite during development of 2-21gal4/UAS-mCD8::GFP, mhc1 embryos from ~14h to 15h AEL (Movie2). (b) Image of neuron at ~15h AEL co-imaged with Ecad::tomato. Inset: Close up of primary dendrite following Ecadherin cell boundaries as indicated by the white arrowheads. (c) Schematic: Control and ablated neurons part of the same embryo. Red cross indicates site of ablation. Top Panel: Dendrite development of Control neuron (Movie3). Bottom panel: Dendrite development of ablated neuron (Movie4). Dashed line indicates point of ablation throughout development. Scale bars = 10μm

When E-cadherin was co-expressed with the 2-21gal4 line to label the adherens junctions of overlying epithelial cells, the vpda primary dendrite coincided with the cadherin signal in 70% of the cases (17/24) (Fig. 2b). Interestingly, even when the primary dendrite did not coincide with cadherin signal, it tracked parallel to these cell boundaries (Fig. S2). This hints at the presence of a cue present in this region that guides the primary dendrite.

To assess the robustness of primary dendrite development, we mechanically perturbed growth by focusing an infrared laser to a point on the primary dendrite to ablate or cut a part of it at ~16h AEL and subsequently observed its recovery. We hypothesized that the dendrite would not be able to maintain growing dorsally as the guidance cue may be lost, since the ablation was carried out at a time point after the completion of primary dendrite development. However, in all examined cases (n=15), ablated primary dendrites recovered from the ablation and could re-grow and continued to extend dorsally (Fig. 2c, Movie3, Movie4). This observation suggested that a persistent cue guided the primary dendrite. Together, these observations support the idea that the primary dendrite number and direction of growth are defined by deterministic rules.

### Morphogenesis of the secondary dendrites is stochastic

Secondary branch growth was much more dynamic than that of the primary branch (Fig. 1). There was also a higher variation in terms of numbers and growth direction in secondary branches. These observations argued against a strictly deterministic mode of growth and we therefore further investigated secondary dendrite growth properties.

Our analysis at the four developmental time points showed a high variability in secondary branch number. For example, at L3, it ranged from 10 to 25 secondary branches per neuron (mean 17± 3)(Fig. 1b). This is consistent with stochastic systems where the features of branching are quite variable but are distributed around a peak when analyzed statistically. However, this observed variation might reflect intersegment variability as neurons from all the abdominal hemi-segments were analyzed. Thus, we pooled data from dendrite arbors from the same abdominal segment (A3) of larvae at 96h ±3 AEL (Fig. 3a). The distribution of secondary branch number ranged from 11-24 secondary branches with a peak at 18±2.6 (Fig. 3b), similar to our previous analysis.

**Figure 3.**
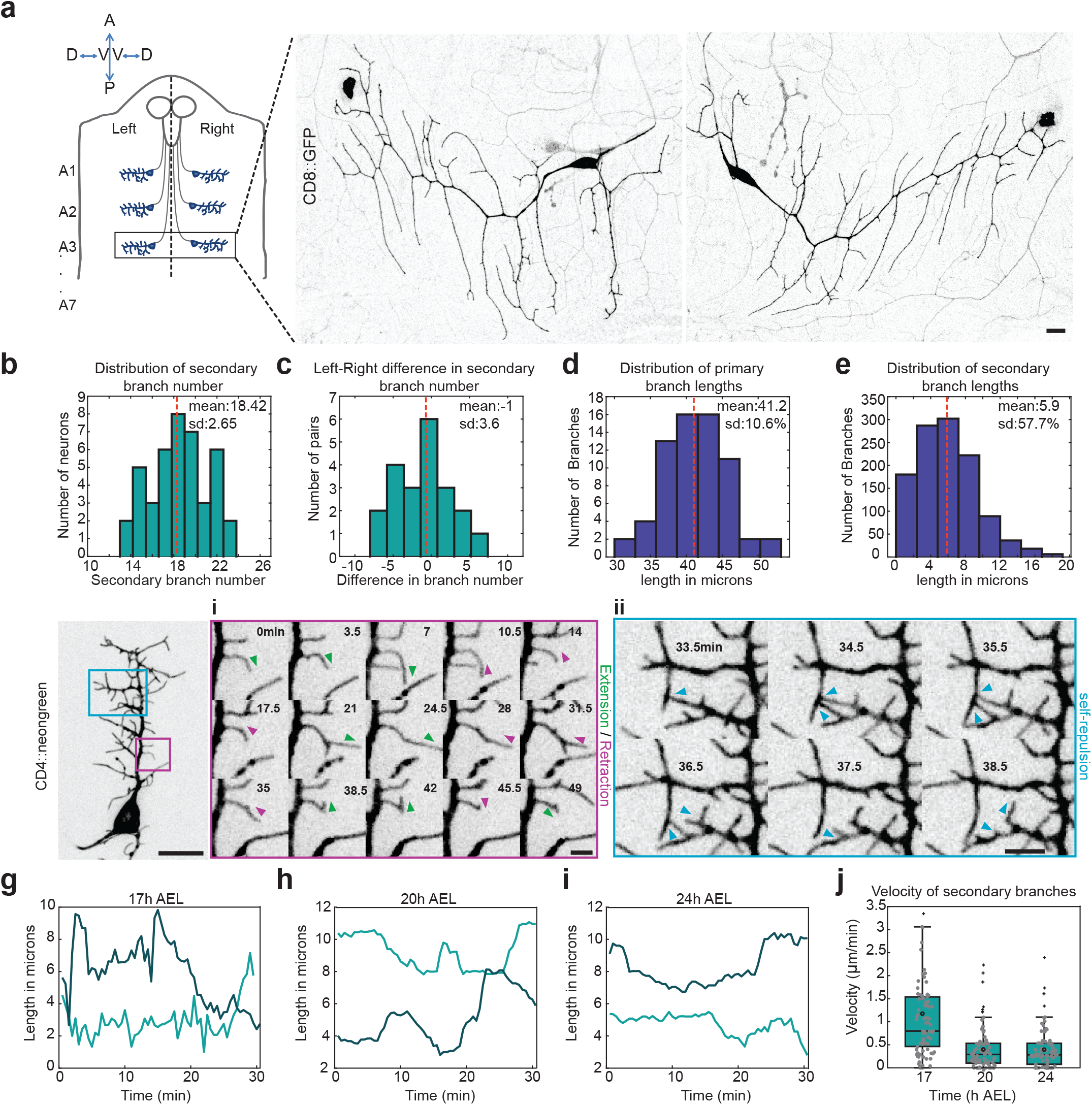
The development of the secondary dendrites is consistent with stochastic growth. (a) Schematic: Larval filet (indicating vpda neurons in the abdominal hemi segments. Neurons in segment A3 were used for analysis. Images of A3 Left and Right vpda neurons at 96h AEL ±3 from homozygous 2-21gal4/UAS-mCD8::GFP larvae. The panel has the following orientation: Anterior at the top, Posterior at the bottom, ventral is the center and dorsal to the sides. (b) Distribution of the total number of Secondary Branches n=42 (21 left-right pairs). (c) Distribution of difference in number of Secondary branches between the left-right pairs. (d) Distribution of primary branch lengths n=66 dendrites from 66 neurons (e) Distribution of Secondary branch lengths analyzed from homozygous 2-21gal4/UAS-mCD8::GFP embryos at 22h AEL ±2 n=1140 branches from 66 neurons (f) Neuron at ~18hAEL.Time-lapse of secondary branch development (Movie5). (i) Green arrowheads indicate extension of a single secondary extension and magenta arrowheads indicated retraction of the same secondary extension. (ii) Blue arrowheads indicate examples of self-repulsion. Panel orientation: dorsal at the top, anterior to the left. (g-i) Lengths of secondary extensions over time, at 17h (g) 20h (h) and 24h AEL (i). The two trend lines indicate two individual branches at each time point. (j) Velocities of secondary extensions at 17, 20 and 24hAEL. Scale bars = 10μm.

However, we could not rule out the possibility that this observed variation might be an outcome of the variability in the age of the larvae. Hence, we compared dendrites from the left and right A3 hemi-segment of the same larva. The difference in secondary branch number between the left-right pairs (Fig. 3c) ranged from −7 to 7 with a mean difference of −1±3.6 between the pairs indicating a high variability of branch number. Thus, the observed left-right asymmetry showed that patterning was not tightly controlled but rather the product of stochastic differences arising during morphogenesis.

Interestingly, the variability of secondary branches between A2 and A3 abdominal segments on the left hemi segments had similar distributions to left-right differences (Fig. S3 a-c), further supporting that intersegment variation is mainly due to stochastic processes.

Further, we measured the lengths of the primary and secondary branches of neurons at ~24h AEL and observed that the variability in primary branch length was 10.6% of the mean length (Fig. 3d) while secondary branches varied by 57.7% of their mean length (Fig. 3e), further supporting the idea that primary branch growth is deterministic while secondary branch development is regulated stochastically.

We next imaged the growth process at a higher temporal resolution (every 30sec) from 17-25hrs AEL (Movie5). In contrast to the unidirectional growth of primary dendrites (Fig. 2a), secondary and tertiary extensions alternated between phases of extension and retraction and, in doing so frequently changed direction (Fig. 3f (i), Fig. 3g-I, each line represents one dendrite). Further, secondary and tertiary extensions retracted upon contact with neighboring extensions, (Fig. 3f (ii)); or self-repulsed. We also observed that the growth rates of the secondary extensions progressively reduced in time. A marked reduction was observed between 17h and 20h AEL. (Fig. 3j).

In summary, secondary dendritic branches are highly dynamic, follow stochastic rules of extension and retraction. A subset of branches is selected from an initial higher number of extensions by self-repulsion. This led us to explore how branching patterns emerge from the stochastic nature of branch dynamics using a computational model.

### A computational model of dendrite branching

The typical class I dendrite morphology is characterized by both a relatively low branch density (with a total dendrite length per unit area of 0.0104 /μm) and smaller dendritic fields as compared to the other morphological classes such as class IV neurons (where, for instance, the total dendrite length per unit area is 0.07/μm)^54^. Inspired by these observations, and with the aim of identifying key mechanisms responsible for patterning this structure, we developed a computational model for stochastic branching morphogenesis of the secondary and tertiary dendrites during embryogenesis.

In our two-dimensional model (Fig. 4a), the extensions were modeled as polymers that grow (extend) and shrink (retract) (see Methods). We considered an initial condition of a fully formed primary branch of length *L*_1_ = 30μm, which remained unaltered during the simulation. This situation corresponds to the state of the neuron at ~15h AEL *in vivo*. New secondary extensions are created from the side of the primary branch at a rate *λ*_1_. At a simulation time corresponding to *t* = 16h AEL, tertiary (resp. quaternary) extensions are also created from the sides of pre-existing secondary (resp. tertiary) extensions at rates *λ*_2_ (resp. *λ*_3_). Once an extension appears, it is either in a polymerizing (i.e. growing) or depolymerizing (i.e. shrinking) state with growth or shrink velocities *ν*_(2,3,4),*on*_ and *ν*_(2,3,4),*off*_, where the subscript number stands for the order of complexity of the extension, and a persistence length *l_p_*. An extension can spontaneously change from a polymerizing state to a depolymerizing one (or *vice versa*) with a rate *k_off_* (or *k_on_* respectively). To complete this model, taken from *in vivo* observations (Fig. 3f(*ii*)) we also introduced a contact-induced depolymerization, that results in what we refer to as ‘self-repulsion’: when an extension encounters its neighbor during polymerization, it will stop growing and shrink with probability *p_off_*. Unless otherwise stated, we set *p_off_* = 1, which implies that the extension will always retract after contacting another extension.

**Figure 4.**
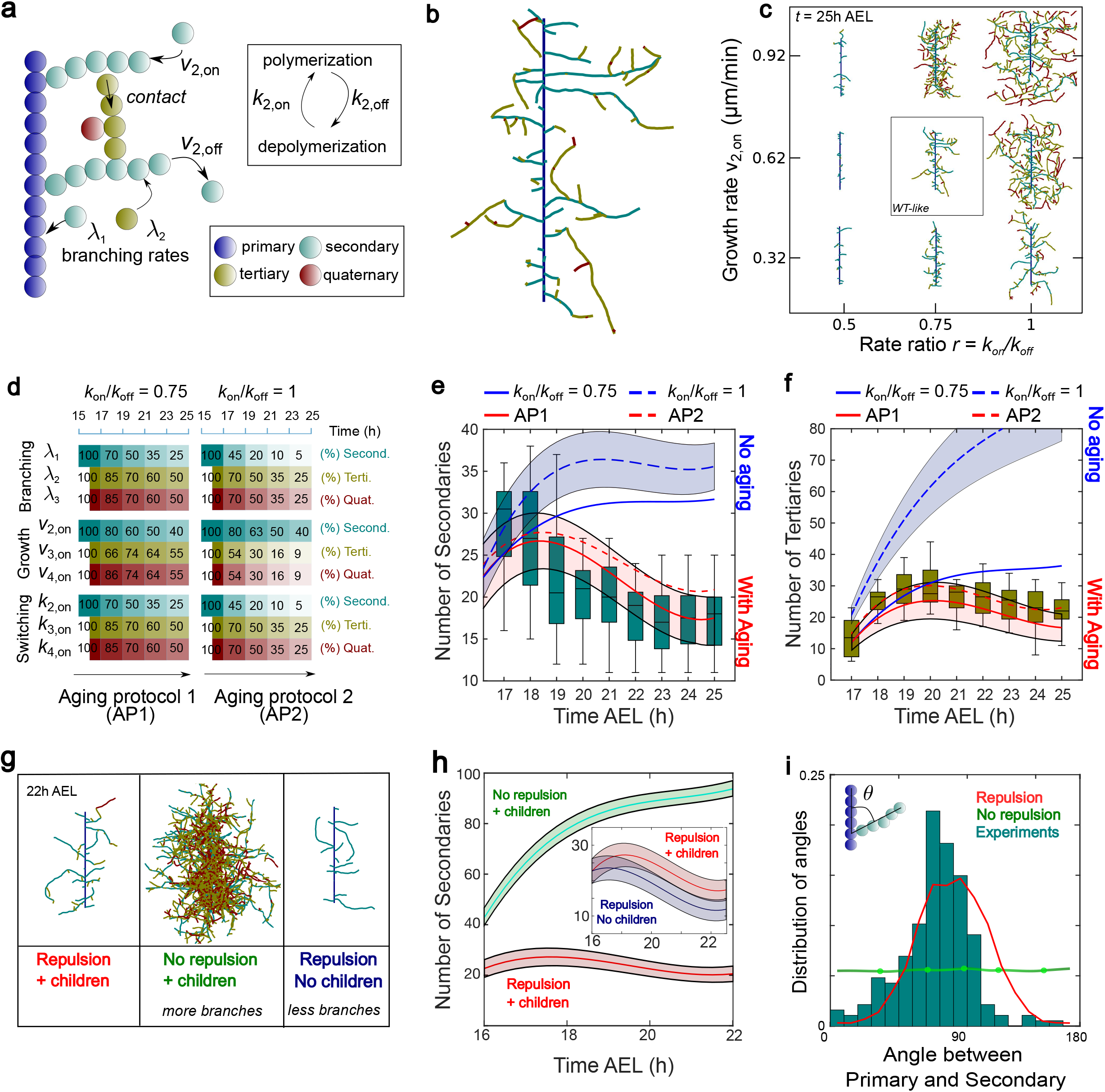
A computational model of dendritic branching and feedback of tree architecture on secondary branch stabilization and orientation. (a) Schematic representation of the mechanisms described in the computational model. (b) Snapshot of a dendritic architecture at 25h AEL from one simulation. (c) Diagram of dendritic morphologies at 25h AEL for different values of the ratio of spontaneous switching rates *r* = *k_on_*/*k_off_* and growth rate of secondary branches *ν*_2,*on*_. The growth rates of tertiary and quaternary branches are given by *ν*_3/4,*on*_ = *ν*_2,*on*_/2 and the values of the other computational parameters are fixed and detailed in Table 1, Option 1. (d) Scheme of the aging protocols AP1 and AP2 implemented for *r* = 0.75 and *r* = 1, respectively. The reduction were performed every 2h and the scheme shows the resulting percentage of the initial parameter value after the reduction at each interval. The depolymerization and switching rates *ν_i,off_* and *k_i,off_* are reduced using the same protocol that for *ν_i,on_* and *k_i,on_*, respectively. (e) Number of secondary branches and (f) Number of tertiary branches over time from 16-25h AEL. Both in (e) and (f), blue lines represent simulations without aging implementation, red lines represent simulations with the two different aging protocols and the boxes represent the experimental data. (g) Snapshots of the dendritic structures at 22h AEL for three different situations: with self-repulsion and child branches, existence of child branches but suppression of self-avoidance and with self-repulsion but without child extensions. (h) Main: Number of secondary branches over time from 16-22h AEL for the models with self-repulsion and child branches (red line) and with child extensions but no self-avoidance (green line) using AP1. Inset: Number of secondary branches over time from 16-23h AEL for the models with self-repulsion and child branches (red line) and with self-avoidance but no child extensions (blue line) using AP1. (i) Probability distribution of the angles between primary and secondary branches, measuring at the end point, at 22h AEL. The histograms represent *in vivo* observations while red and green lines represent simulations with and without self-repulsion interactions using AP1. Similar results than the one showed in (h) and (i) for AP2 can be found in Fig. S4i-j. The number of simulations for blue lines in (e) and (f) was *N_sim_* = 50 and for red lines in (e) and (f), and all cases in (h) and (i) was *N_sim_* = 500. In all figures, simulation lines represent the average value of the realizations and shadows represent the 1*σ* confidence interval. In addition, a 2-point time-averaging was performed using spline interpolation for lines in (e), (f) and (h). For *r* = 0.75 and AP1 in (e) and (f), and all cases in (g), (h) and (i) the initial values of the computational parameters are detailed in Table 1, Option 1. For *r* = 1 and AP2 in (e) and (f) the initial values of the computational parameters are detailed in Table 1, Option 2. For each box corresponding to *in vivo* observations, the central line is the median, the box extends vertically between the 25th and 75th percentiles and the whiskers extend to the most extreme data that are not considered outliers.

We first explored the space of constitutive parameters of the model and found that a range of different dendrite morphologies can be described using this computational scheme. In particular, we focused on three sets of parameters that played a fundamental role in dendrite architecture: branching rate (*λ*), growth rate (*ν*) and the ratio of spontaneous switching rates, *r* = *k_on_*/*k_off_*. We found that growth velocity tuned the branch length as well as the total area of the dendrite: the higher the velocity, the larger the average length of branches and area (Fig. 4c). On the other hand, branching rate *λ* impacted branch number, such that, as it increased the total number of branches also increased (Fig. S4a,d). Importantly, we discovered that the ratio of switching rates *r* was the most relevant parameter to explore the different types of emerging structures. For any value of both branching rate and growth velocity we could account for the wide phenomenology described by this model by only changing the value of *r* (Fig. 4b-c). For low values of *r* (i.e. a higher probability of switching from a polymerizing to a depolymerizing state) only a few short secondary extensions survived to the final stages. For intermediate values of *r*, a moderate branch density emerged, similar to class I dendrites. Finally, for values of *r* close or equal to 1, we obtained structures with high branching density and long branches, leading to final morphologies with very large total areas (Fig. 4c). This qualitative trend was observed irrespective of both growth and branching rates (Fig. 4c, S4a-b).

**Table 1.** Computational parameters.

| Parameters |  | Notation | Option 1 | Option 2 |
| --- | --- | --- | --- | --- |
| Time step (**min**) | | $\Delta t$ | 0.5 | 0.5 |
| Length primary branch (**μm**) | | $L_1$ | 30 | 30 |
| Number of beads in primary branch | | $N_1$ | 100 | 100 |
| Branching rate<br>(**μm<sup>-1</sup> min<sup>-1</sup>**) | Secondary | $\lambda_1$ | 0.22 | 0.12 |
| | Tertiary | $\lambda_2$ | 0.05 | 0.05 |
| | Quaternary | $\lambda_3$ | 0.02 | 0.02 |
| Growth/shrink rates<br>(**μm min<sup>-1</sup>**) | Secondary | $v_{2,on/off}$ | 0.62 | 0.32 |
| | Tertiary | $v_{3,on/off}$ | 0.31 | 0.18 |
| | Quaternary | $v_{4,on/off}$ | 0.31 | 0.18 |
| Persistence length (**μm**) | | $l_p$ | 17 | 17 |
| Spontaneous switching rate to a polymerizing state<br>(**min<sup>-1</sup>**) | | $k_{on}$ | 0.5 | 0.67 |
| Spontaneous switching rate to a depolymerizing state<br>(**min<sup>-1</sup>**) | | $k_{off}$ | 0.67 | 0.67 |

However, despite an extensive parameter exploration, we were unable to find quantitative agreements between *in vivo* and simulated branches; the time evolution of the number of branches simulated branches was consistently above *in vivo* values (Fig. S4c-d, Fig. 4e-f (blue lines)). Motivated by the *in vivo* observation that the dynamics of secondary extensions slows down over time (Fig. 3g-j), we incorporated a progressive decay of all dynamic parameters in our computational model. We implemented an “aging factor” by reducing the rates of growth, branching and switching with time. By varying both intensity and kinetics of aging, we obtained different types of dendrite arborization patterns that ranged from barely affected highly active dendrites (for small aging factors) to dendrites that nearly froze at early stages exhibiting low density arborization patterns (for high aging factors). Additionally, the larger the ratio between switching rates was, the more intense the aging factor needed to be in order to obtain adequate numbers and branch lengths at intermediate and final time points consistent with experiments (Fig. 4d-f). To explore this, we chose two sets of parameters, with different values of the switching ratio *r*, and ran simulations by applying two different aging protocols; AP1 with *r* = 0.75 and AP2 with *r* = 1 (Fig. 4d). The resulting branch numbers and lengths were in good agreement with *in vivo* observations for both protocols (Fig. 4e-f (red lines) and Fig. S4f-g). Notably, for one of the parameter sets (AP1 in Fig. 4d) the value of the growth velocity was set to 0.62μm/min, which is in accordance with *in vivo* measurements (Fig. 3j).

Thus, we developed a computational model that predicts the emergence of a range of possible dendrite morphologies from a few local statistical features of branch dynamics. For a wide subset of the parameter space, the combination of all the different elements of the model lead to arborization patterns consistent with *in vivo* observations (Fig. 4d-f, S4f-g and Movie6).

### Feedback of tree architecture on secondary branch stabilization and orientation

While the dynamics of branches is partly governed by a few constitutive parameters that are homogeneous in space, simulations also revealed the importance of feedback of the structure of the dendritic tree onto local branch dynamics.

In our movies we observed that when a secondary extension (parent branch) depolymerizes, it does not shrink beyond the branching point of any existing tertiary extensions (child branches) (Fig. S3d, Movie7). Our simulations implemented this observation by imposing a rule which prevents the depolymerization of a parent branch beyond the forking point with its child branch. Instead, the branch grows again from the forking point. With this mechanism, when a parent branch depolymerizes, it is protected from complete shrinkage by the existence of child-branches. Consistently, in simulations, we observed fewer secondary branches when tertiary and quaternary extensions (children) were not allowed to develop (Fig. 4g-h (blue lines)). Thus, the proliferation of tertiary and quaternary extensions reduces the probability of complete branch elimination giving rise to branch self-stabilization.

The most striking feature of secondary branch development was contact-induced depolymerization or self-repulsion, which occurred quite frequently. This hinted at an important role for self-repulsion in dendrite patterning and hence we decided to explore this feature further.

When a new branch is created in a region surrounded by other branches, its probability of survival strongly decreases as its probability of encountering other branches increases as a result of contact-induced depolymerization. Thus, the effective densities of branches feeds back negatively on branch number (Fig. 4g,h (red lines)), an effect that has a deeper impact on tertiary branches, which have an increased probability of encountering neighboring branches (Fig. S4h (Top)). Additionally, branch lengths are also constrained by this mechanism (Fig. S4h (Bottom)). These two phenomena were confirmed in simulations of both, dendrites with and without self-repulsion (Fig. 4g,h (red and green lines respectively)). Therefore, self-repulsion strongly reduces both, branch length and number, the defining feature of class I dendrites.

Self-repulsion also had unexpected impact on dendrite orientation. We measured the distribution of angles made by the secondary dendrites on primary branches both in simulations and experiments. We obtained a peak around 90° while lower frequency values were found around both 0° and 180°. Thus, secondary dendrites tend to be oriented orthogonally with respect to the primary branch (Fig. 4i). To test if self-repulsion played a role in orienting branches, we performed a simulation in which contact-induced depolymerization was eliminated and found that it led to a uniform angle distribution (Fig. 4i (green line)). Hence, in the simulations of vpda neuron development, the observed orientation of secondary dendrites does not require a guidance cue per se, but is an outcome of a selection process for branches that minimizes branch density and cross-over by contact-induced depolymerization or self-repulsion. Thus, self-repulsion is potentially a key feature responsible for the characteristic shape of class I vpda neurons *in vivo*.

In summary, in our model, the geometry of the dendritic tree that emerges from the local statistical features of branch dynamics exerts positive (self-stabilization by child-branches) and negative feedbacks (branch elimination and shrinkage due to contact-induced depolymerization) on branch dynamics by effectively changing the rate of shrinkage.

### Dscam1 restricts the orientation of secondary dendrites

These results led us to further investigate the potential impact of self-repulsion in secondary branch organization *in vivo* using *dscam1* mutants. Dscam1 is a type I membrane protein of the immunoglobulin superfamily ^55^ expressed in all 4 neuronal classes. Alternative splicing can potentially generate more than 38,000 Dscam1 isoforms such that every cell expresses a different isoform, which defines its identity. Dscam1 is expressed in the multi-dendritic neurons and its loss causes self-avoidance defects in all 4 classes of md-neurons^56–58^ (see also Fig. 5a). All previous studies analyzed the effect of Dscam1 at a stage when dendrite patterning was already complete. However, how self-avoidance precisely affects the emergence of dendrite pattern during morphogenesis remains unknown.

**Figure 5.**
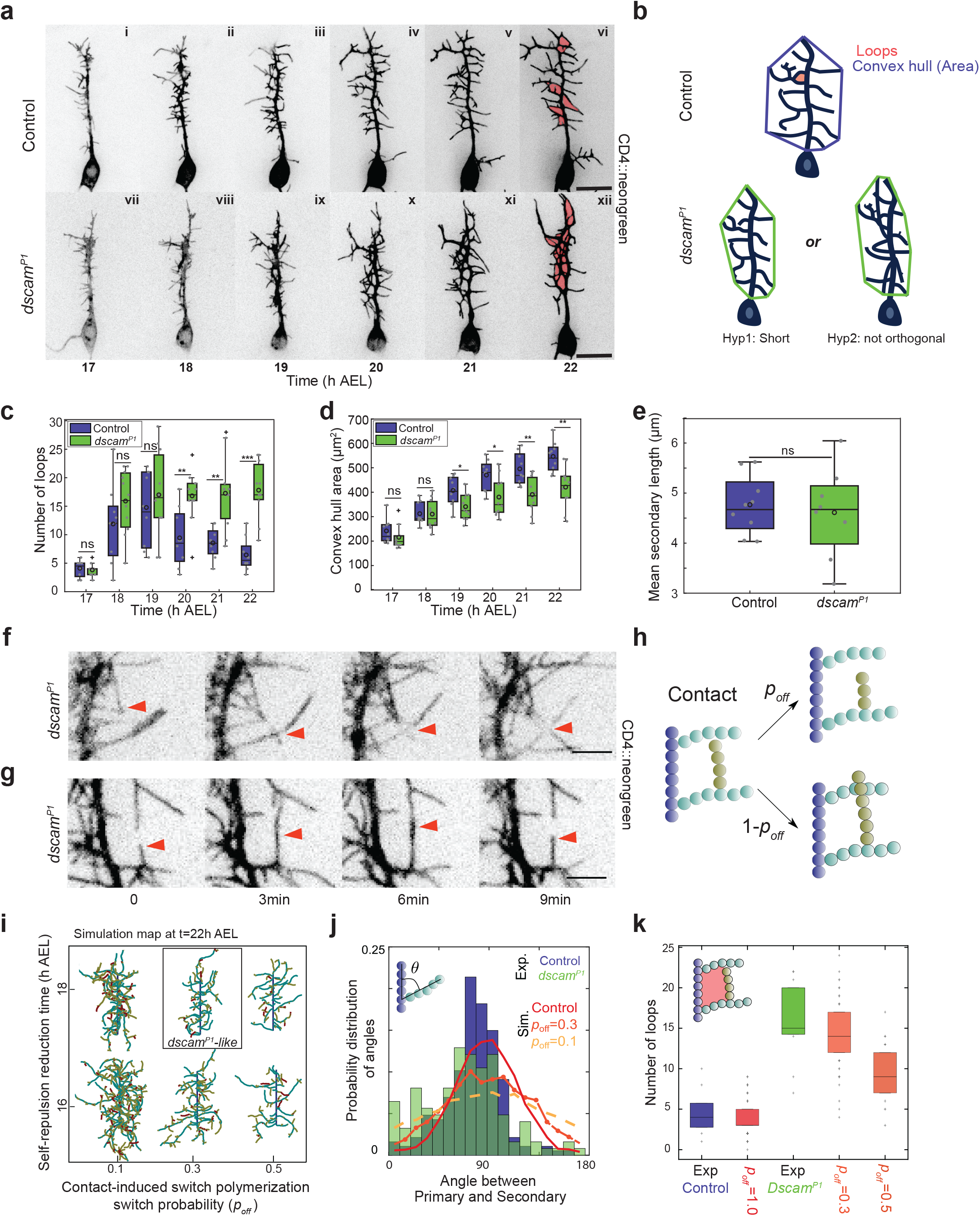
Dscam1 dependent self-repulsion restricts orientation of dendrite development. (a) Top panel: Time-lapse images of dendrite development of 2-21gal4/UAS-mCD4::neongreen (Movie8) Lower panel: Time-lapse images of dendrite development of *dscam^P1^*/ΔDscam1; 2-21gal4/UAS-mCD4::neongreen (Movie9). The embryos were injected with Ryanodine. Imaged from ~17h AEL to ~22h AEL.(b) Schematic: Area is calculated as the area of the convex hull that contains the dendrites. Red shaded area indicates the loops formed by dendrite crossovers. Top panel: Control dendrites Bottom panel: *dscam^P1^* dendrites could have a smaller area because of shorter dendrites (ii) or dendrites that are not perpendicular.(c) Number of loops in the dendritic tree ~17-22h AEL. (d) Area of the dendritic field covered by the dendrites ~17-22h AEL. .(e) Quantification of mean secondary branch lengths ~17-22h AEL. For (c) and (d) and (e) Control (blue) n=9 Mutant (green) n=11 (f)Loss of repulsion: Red arrows indicate a dendrite crosses over another branch and does not retract. (g) Remnant repulsion: Red arrows indicate an example of remnant self-repulsion *dscam^P1^* (Movie10). (h) Schematic: Upon contact with a neighbor, an extension either crosses over its neighbor with a probability of 1 − *p_off_* or switches to a depolymerizing state with a probability *p_off_*. (i) Diagram of dendritic morphologies obtained in simulations at 22h AEL for different values of *p_off_* introduced at different developmental time points. (j) Number of loops at 22h AEL for Control experiments, simulations with self-repulsion (*p_off_* = 1), *dscam^P1^*, *p_off_* = 0.3, *p_off_* = 0.5. The number of simulations for all cases was *N_sim_* = 50. (k) Probability distribution of angles between primary and secondary branches Control(blue histogram) n=187, Mutant(green histogram), n=124 ks test *p* = 0.0113. Simulation control (red), *p_off_* = 0.3 (orange), *p_off_* = 0.1 (yellow). Simulation lines represent the average over *N_sim_* = 100. For each box, the central line is the median, the ‘o’ is the mean, the box extends vertically between the 25th and 75th percentiles, the whiskers extend to the most extreme data that are not considered outliers, and the outliers are plotted individually. Statistical significance for (c), (d) & (e) has been calculated using Mann-Whitney U test. ns, * p<0.05; ** p<0.01, *** p<0.001. All the panels have the same orientation: dorsal at the top, anterior to the left. Scale bars = 10μm. In all the simulations, the implemented aging protocol is AP1 (Fig. 4d) and the initial values of the computational parameters are detailed in Table 1, Option 1.

The embryonic development of vpda neurons was observed in *dscam^P1^* mutant^55,56^ over a *dscam1* deletion line, marked with CD4::neon green. The mutant and control embryos were imaged from 17h to 22hAEL (Fig. 5a, Movie8 & 9). The development of the primary branch was unaffected in the mutant (Fig. S5a), with no significant difference in the primary branch length measured at 22hrs AEL (Fig. S5b).

At a first glance, at 22h AEL the *dscam^P1^* dendritic tree does not appear to share the high branching density of the simulations with no self-repulsion (Fig. 4g (middle panel)). We carefully examined the development *dscam1* mutant dendrites in order to how *dscam^P1^* is different from the no-repulsion simulations.

Consistent with previous studies performed at later stages, at 22h AEL, mutant dendrites frequently crossed over each other (Fig. 5f), thereby forming apparent ‘loops’ in the dendrite architecture (Fig. 5a (xii) & Schematic 5b). These ‘loops’ were also observed to a lesser extent in control dendrites (Fig. 5a (vi)). The number of loops increased in both control and mutant embryos from 17 to 19h AEL. However, from 20-22h AEL, the number dropped strongly in control dendrites, but was maintained in *dscam1* mutants (Fig. 5c). This implies that from 17h to 19h AEL, the secondary and tertiary dendrite extensions explore the space around them and contact their neighbors in both genetic conditions. However, these contacts are not maintained in control embryos due to contact-induced repulsion but persist in the *dscam1* mutant.

We also observed that the surface area of the dendritic field covered by the dendrite in mutants was smaller than in controls (Fig. 5d). The area of the convex hull (smallest polygon containing the dendrites) of the dendrites (Fig. 5b) became significantly different between controls and *dscam1* mutants from 19h AEL and was 20% smaller in the *dscam1* mutants at 22h AEL (Fig. 5d). The smaller dendritic field areas observed in the mutants could either reflect that the secondary branches are shorter (Fig. 5b Schematic (Hyp1)) or that the secondary branches are not perpendicular to the primary branch (Fig. 5b Schematic (Hyp2)) as shown in simulations without self-repulsion (Fig. 4i). To test these hypotheses, we measured secondary branch lengths at 22h AEL and found no significant difference between mutants and controls (Fig. 5e). However, we observed a significant reduction in the number of perpendicular branches in *dscam1* mutants (Fig. 5j). This suggested that the reduced dendritic field is due to a loss of perpendicular alignment of secondary branches. However, the angle distribution of secondary branches was flatter but not uniform as predicted by the simulations with ‘no-repulsion’, suggesting that self-repulsion is not completely lost in *dscam1* mutants. Upon close inspection, indeed, self-repulsion was not completely blocked in *dscam1* mutants (Fig. 5g, Movie 10).

To gain further quantitative assessment of self-repulsion in *dscam1* mutants, we compared these experimental observations with simulations. We therefore modified our simulations of no-repulsion such that during dendrite development, the probability of switching to a depolymerizing state *p_off_* was reduced upon contact with neighboring dendrites (Fig. 5h). Using this modification we tested a range of values for *p_off_* while for control simulations we maintained *p_off_* = 1. For *p_off_* = 0.1 (implying a 10% probability of retraction after contact) very dense arborization patterns were obtained, (Fig. 4g) and increasing the value of *p_off_* was accompanied by a decrease in branching density (Fig. 5i). In particular for *p_off_* = 0.5 the final dendritic morphology exhibits a low branch density and a tree-architecture similar to control cells *in vivo* (Fig. 5i). In addition, we observed highly dense arborization patterns at 22h AEL when *p_off_* was reduced at 16h AEL (for *p_off_* = 0.1 and *p_off_* = 0.3). However, when *p_off_* was reduced at 18hAEL the arborization patterns obtained were similar to *in vivo* observations. Hence, focused on an implementation time of 18h AEL.

Further, we measured both the distribution of angles between primary and secondary branches. For *p_off_* = 0.1 the distribution becomes flatter, implying a higher reduction of perpendicular branches in comparison to *dscam^P1^* (Fig. 5j). However, *p_off_* = 0.3, the angle distribution was consistent with the *dscam^P1^* mutant data. We then measured the number of loops at 22h AEL. For *p_off_* = 0.3 the number of loops is consistent with *dscam^P1^* data (Fig. 5k) while for *p_off_* = 0.5 the dendrite configurations have significantly fewer loops than in *dscam^P1^*. Thus, our simulations also support the experimental observation that self-repulsion is reduced but not fully suppressed in *dscam^P1^* mutants.

To conclude, self-repulsion patterns secondary branches by restricting direction of secondary extension growth, thus, ensuring a preferential stabilization of secondary branches perpendicular to the primary branch, giving the class I neurons their characteristic shape.

## Discussion

Several studies over the last two decades have identified and characterized molecules important for regulating different aspects of pattern formation of dendrites ^2,53,59–62^. How a neuron integrates molecular information to generate its characteristic dendritic shape or the ‘rules of branching’ followed by developing dendrites is still unclear. Addressing this type of question requires observing dendrite morphogenesis live and *in vivo*.

The embryonic development of the *Drosophila* sensory neurons has been described before ^52,53^. These studies were important to provide a description of when and how the primary and secondary branches were initiated and when patterning was complete. However, a precise quantitative analysis of development was necessary to deepen our understanding of this process. Our study quantitatively described the embryonic development of the class I vpda neuron and showed that the primary dendrite grows deterministically while secondary dendrite growth follows stochastic rules.

The primary dendrite followed the cell outlines of the overlying epithelial cells (Fig. 2c). This suggested that the primary dendrite might be guided by an extrinsically patterned cue present in this region, eg. at cell-cell interfaces. Signals classically involved in axon guidance like Semaphorins ^63,64^, Slits ^22,24^ and Netrins ^25^ have also been implicated in dendrite targeting. Sax-7 is a homologue of the mammalian L1 cell adhesion molecule is extrinsically patterned in the *C.elegans* hypodermal cells and guides dendrite patterning of the PVD neuron. It is therefore very likely that such a molecule guides the primary dendrite of the vpda neuron as well ^28^. Interestingly, the cell boundaries along which the primary dendrite grows appear to be stretched as junctions are all aligned along the dorsal-ventral axis and MyosinII is enriched along these boundaries ^65^. Thus, alternatively cortical tension might be involved in guiding the primary dorsal branch. Tension could for instance guide the primary dendrite by forming a path of least resistance for growth through local tissue deformation or make a stiffer substratum for dendrite growth ^66^

The development of secondary extensions was studied both *in vivo* and *in silico* using a computational model relying on stochastic rules of growth and branching dynamics, with self-avoidance interactions. The onset of secondary branch morphogenesis was very dynamic and as morphogenesis progressed, the dynamics greatly reduced. After 22h AEL the number and lengths of secondary branches was stable. The computational model demonstrated the necessity to incorporate the decay of its kinetic parameters (‘cell aging’) to account for bounded growth of dendrites. It is possible that dendrites age through stabilization by microtubules ^59,67^. Tagged actin was expressed in the highly dynamic dendritic tips while microtubules were less dynamic and present in the more stable parts of the dendritic tree (data not shown). Thus, aging could require a mechanism that decreases actin dynamics and increases microtubule polymerization over time.

This work reveals the importance of two opposing phenomena of tree architecture feedback on local branch dynamics. “Child-branches” (tertiary) have a stabilizing effect on parent branches (secondary). Thus, branches with more children have a higher survival probability. However, a high density of branches exerts also a negative feedback due to contact-induced depolymerization. The most striking feature of secondary branch development is self-repulsion, which occurred very frequently within populations of growing secondary and tertiary extensions. Self-repulsion increases the branch shrinking probability introducing a new effective depolymerization rate *k*′_*off*_, which is proportional the product of the probability of encounter with another branch and the probability of changing to a depolymerizing state when the encounter takes place, *p_off_*. Remarkably, while all the previously described computational parameters are constitutive quantities of the model, *k*′_*off*_ is a variable parameter that depends, among others, on the local branch density. Through this ‘geometric’ feedback, the tree architecture updates dynamic parameters of the dendritic tree and affects its own morphogenesis. Thus self-stabilization and self-repulsion constrain the number and lengths of the secondary branches.

Our study revealed another remarkable function of self-repulsion in biasing the orientation of secondary dendrites perpendicular to the primary branch. The prediction from simulations was confirmed *in vivo* through the analysis of *dscam1* mutants. In *dscam1* mutants, self-repulsion is reduced so that, in most cases, branch growth persists after self-contact in all directions (Fig. 5j). Thus for a given dendrite, its final orientation is dependent on where and when it encountered sister dendrites. Hence, both experimental observations and simulation argue that the dendritic tree-patterns are not guided deterministically, but emerge as self-organized structures from the stochastic nature of secondary branch development.

In contrast to the prediction of the “no-repulsion” simulations, the mean lengths of the *dscam^P1^* branches were not significantly longer than control branches. However, the lengths of the non-orthogonal branches were twice as long as the orthogonally oriented branches in *dscam^P1^* condition (Fig. S5c) supporting that self-repulsion reduces branch length. Also in contrast to the “no-repulsion” simulations, the number of secondary branches in the *dscam^P1^* mutants was significantly lower than in control dendrites (Fig S5d). Thus, self-repulsion does not account for all the function of Dscam1. Dscam1 is a complex molecule that could have several unknown functions. One possibility is that its loss might impinge on a signaling pathway necessary for stabilization of secondary extensions.

While our data reveal self-organizing properties of dendrites, our observations do not rule out the possibility that extrinsically patterned cues are also involved in secondary branch morphogenesis. We suggest that self-organizing principles reported in this study provide a ‘ground state’ upon which additional cues could operate. For example, the anterior-posterior asymmetry in secondary branch length of the vpda neuron (data not shown) depends on extrinsically patterned ten-m ^68^, albeit, at later stages. In the class III neurons, Netrins are necessary to target at least one dendrite to the lateral chordotonal organ, but Dscam1 plays a major role in regulating the even dendritic field coverage ^25^. The role of extrinsic cues would have to be analyzed in the background of stochastic branching processes and geometric feedbacks, which, by themselves provide spatial bias to the growth of secondary and tertiary dendrites.

This study highlights the dynamics of dendrite growth and provides the analytical basis for further investigating the dynamics of dendrite growth and patterning *in vivo*. It will be particularly interesting to explore, on the basis of the present study, how class specific morphogenesis of multi-dendritic neurons emerges during development, as a function of varying class-specific self-organizing rules of dendritic arborization. We suggest that such intrinsic rules, though statistical, might be genetically encoded. Ultimately, understanding how genes determine statistical rules of branching provides an opportunity to understand how, in a broader context, genes encode form.

## AUTHOR CONTRIBUTIONS

A.P. and T.L. conceived the project and planned experiments. A.P. performed all experiments, quantifications and data analysis. N.T.-E. and J.-F.R. designed the model and the computational simulations; N.T.-E. obtained the numerical results. All authors analyzed the results and wrote the manuscript.

## ACKNOWLEDGEMENTS

We thank all members of the Lecuit lab for very useful discussions throughout the course of this work, for creating a stimulating environment, and also for critical comments on the manuscript. We thank members of A.P.’s thesis jury, Markus Affolter, Edouard Hannezo and Peter Soba for fruitful and insightful comments. We thank Jean-Marc Philippe for constructing the neon green fly line and Benoit Aigouy for the ‘Dendrite arborization Tracer’. We are grateful to the IBDM imaging facility for assistance with maintenance of the Microscopes, and FlyBase for maintaining curated database and Bloomington fly facility for providing transgenenic flies. AP. was supported by Ph.D. fellowship from the LabEx INFORM (ANR-11-LABX-0054) and of the A*MIDEX project (ANR-11-IDEX-0001-02), funded by the “Investissements d’Avenir French Government program”. NT-E was supported by the “Investissements d’Avenir French Government program” managed by the French National Research Agency (ANR-16-CONV-0001) and from the Excellence Initiative of Aix-Marseille Univerité - A*MIDEX. We acknowledge France-BioImaging infra-structure supported by the French National Research Agency (ANR-10-INBS-04-01, «Investments for the future»).

## MATERIALS AND METHODS

### Fly strains and genetics

The following fly strains were used; 2-21 gal4 (From Yuh-Nung Jan), mhc 1 mutant ^69^, UAS-CD8::GFP (Bloomington stock #5137), UAS-CD8::RFP (Bloomington stock #27399), Ecad::Tomato (Bloomington stock #58789), UAS-Dscam1GFP (Bloomington stock #66203) ^56^ *dscam^P1^* (Bloomington stock #11412), ΔDscam1(Bloomington stock# 9062).

### Molecular cloning and transgenic fly lines

To generate the UAS-CD4::neongreen flies, pACU2_CD4-mIFP T2A HO1 from Xiaokun Shu ^70^ ordered from ADDGENE (72441) was modified. The open reading frame (ORF) part corresponding to mIFP-T2A-HO1 was replaced by mNeonGreen’s one (a bright modified GFP, Shaner et al, Nat Methods 2013), producing a Cter mNeonGreen tagged version of the human transmembrane protein CD4. Expression plasmid named pACU-CD4-NeonGreen was verified by sequencing (Genewiz) and sent to Bestgene Inc. to perform transgenesis into attP-containing docking site strains attP2 (3L, 68A4). Sequence of the plasmid is available upon request.

### Protocol for embryonic development of the class I vpda neuron

The muscular tissue is established at the stage of observation; hence, muscle contraction prevented the capture of stable and time resolved images. A previous study suggested dendritic development was normal in mutants for muscle myosin, which paralyzes embryos^71^. Therefore, vpda neurons labeled with CD8::GFP (with the 2-21Gal-4 line) were imaged in a myosin heavy chain mutant background ^69^ to prevent muscular contractions. No statistically significant difference in neuron morphology was observed in mhc1 mutant embryos (Fig. S1 e-h), suggesting that mhc1 mutants provide an effective solution to temporally resolve embryonic dendrite morphologies.

When, recombining the mhc[1] mutant allele with the *dscam^P1^* mutant proved to be difficult; in these experiments, Ryanodine (a drug that inhibits muscle contractions by binding to the ryanodine receptor and preventing calcium release into the sarcoplasmic reticulum) was injected into stage 16 embryos. Single images of water and ryanodine injected vpda neurons were captured 10 hours after injection. Analysis revealed no major defects in dendrite morphology, apart from small but significant increase in the number of secondary branches in the ryanodine injected embryos (Fig. S1h-k). As there were no other patterning differences, the ryanodine-injected embryos still offered an effective solution to obtain time resolved images of *dscam^P1^* mutant embryos.

### Embryo preparation and Drug Injections for live imaging

Embryos were prepared as described before ^72^. In brief, embryos were de-chorionated using bleach, for about 1 minute and then washed thoroughly with distilled water. The embryos were then aligned ventro-laterally on a flat piece of agar and then glued to a glass coverslip. These embryos can be submerged in halocarbon oil and can be imaged directly. Alternatively, glued embryos (;*dscam^P1^*/ΔDscam1;2-21gal4, UAS-CD4::neongreen/’’; 2-21gal4 and ;;UAS-CD8::GFP/’’; 2-21gal4) were kept in an airtight box containing Drierite for about 7 min, then covered in halocarbon oil, and then injected with Ryanodine. Ryanodine (from Merck) was injected at a concentration of 50mM in mid stage 16 embryos about 10 minutes prior to imaging using a FemtoJet 4i microinjector by Eppendorf and then imaged at the microscope.

For analysis of dendrite morphology at different stages of development, whole larvae at L1, L2 and L3 were placed in a watch glass with a few mL of 1mM Sodium Azide to paralyze them and then mounted in low melting agarose to be imaged on a lightsheet microscope so that the larvae could be rotated and all the neurons could be easily imaged.

### Larval fillet preparations and Immunostaining

Homozygous 2-21gal4/UAS-mCD8::GFP larvae (96h±3) were filleted using standard protocols and fixed in 4%PFA for 15min and stained with a rabbit anti-GFP antibody (1:500, Invitrogen). Secondary antibodies conjugates with Alexa488, (Invitrogen) were used 1:500. The stained larvae were mounted in VECTASHIELD®.

### Image Acquisition

For live imaging embryos were prepared as described earlier and time-lapse imaging was performed with a dual camera spinning disc (CSU-X1, Yokogawa) Nikon Eclipse Ti inverted microscope (distributed by Roper) using a 100X/N.A 1.4 oil-immersion objective. The system acquires images using the Meta-Morph software and images were taken as z-series of 1.5μm X10 planes spanning 13.5μm and acquired with a frame rate of 3min for 10hours or 30min for 10 hours or 30s for 1hour.

For whole mount imaging, larvae were treated as described before and imaged on a Zeiss Lightsheet Z.1 on a 20X/N.A 0.8. The system acquires images using ZEN software conditions were kept consistent for all 3 stages.

Fixed larval fillets were imaged on a Zeiss LSM 880 on a 25X/N.A 0.8. The system acquires images using the ZEN software and image stacks with spacing of 0.15–0.2 μm were collected and stack focused projections of 7-10 planes analyzed.

### Laser Ablations

Ablations were performed at around 15-16h AEL on an inverted microscope (Eclipse TE 2000-E; Nikon) equipped with a spinning-disc (Ultraview ERS, Perkin Elmer) for fast imaging. Time lapse at a single z-plane was acquired using a ×100 1.4 NA oil immersion objective. Ablations were performed in parallel with image acquisition. Ablation events were obtained by exposing the primary dendrite, for duration of 2–3 ms, to a near-infrared laser (1030 nm) focused in a diffraction-limited spot. Laser power at the back aperture of the objective was ~400mW. Once ablated, the neurons were imaged on the dual camera, spinning disc (CSU-X1, Yokogawa) Nikon Eclipse Ti inverted microscope (distributed by Roper) using a 100X/N.A 1.4 oil-immersion objective as described above. In these experiments, the control neurons were the unablated neurons from the neighboring segments.

### Image Processing and Segmentation

CD8::GFP or CD4::neongreen was used to label the neuronal cell membranes. A custom ImageJ macro integrating the Stack Focuser plugin from M. Umorin was used to project (by maximum intensity projection the z-planes with signal from the neuron. This resulted in sharper cell outlines and better S/N ratio compared with maximal projections. The 2D projected stacks were then segmented using a custom ‘Dendritic Arborization Tracer’ from B. Aigouy.

### Data Analysis

The ‘Dendritic Arborization Tracer’ provides skeletonized images in such a way every segment (connected to other segments through vertices) is assigned a unique identity (Fig. S1b). We then ordered these individual segments using MATLAB (including Curve Fitting Toolbox, Image Processing Toolbox, Statistics and Machine Learning Toolbox). Starting from the cell body we connected segments that made the longest line and assigned it to be the primary branch. Similarly, we then connected all the segments making the longest lines starting from the primary branch to be secondary branches and so on (Fig. S1c). However, our code does account for loops in the dendritic tree. In the control, the dendrites rarely crossed each other, however, in the event of a crossing, we removed the loops (wild type condition). We then performed all our analysis on this processed skeleton of the dendrite. To plot data, we used Matlab (the IoSR box Plot function by Christopher Hummersone).

In the case of *dscam1* mutants, loops in the dendritic tree prevented us from using the software, thus, the movies of development segmented using the ‘Dendritic Arborization Tracer’ but the dendrites were ordered manually by observing the movies of embryonic development.

### Statistics

For all experiments data points from different neurons/embryos from at least 2 independent experiments were pooled. For each box, in the box plots, the central line is the median, the ‘o’ is the mean, the box extends vertically between the 25th and 75th percentiles, the whiskers extend to the most extreme data that are not considered outliers, and the outliers are plotted individually. All the P values are calculated using a two-sided non-parametric Mann–Whitney test (Matlab statistic toolbox) except in Fig. 5i where P values are calculated using the two sample Kolmogorov-Smirnov test and Fig 1(d) (e) and (f) and Fig 4 where P values are calculated using a paired-sample t-test.

### Computational model

To understand the role that the different mechanisms involved in the class I vpda neuron evolution have in its final morphology we developed a two-dimensional stochastic branching model with contact-induced branch depolymerization (or self-repulsion). Therefore, in contrast to other branching models for morphogenesis ^34^, we include the possibility of branch retraction.

In our simulations, the dendrite structure is discretized into elementary spatial units (hereby called beads). At the simulation time *t* = 0 (approximately corresponding to the developmental time *t* = 15h AEL), we consider a pre-existing primary branch composed by a constant number of beads *N*_1_ = 100 (corresponding to *L*_1_ = 30μm) and situated along the y-axis.

#### Branching

At each time step Δt, a new secondary branch is created from a non-branched bead with a probability denoted by

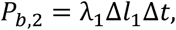

where Δ*l*_1_ is the inter-spacing between beads in the primary branch; λ_1_ is the branching rate of secondary branches. After 1h of evolution (corresponding to t = 16h AEL), tertiary and quaternary branches are allowed to be created using the same mechanism. Similarly, we define the per-bead new tertiary (resp. quaternary) branch creation probability *P*_*b*,3_ = λ_2_Δ*l*_2_Δ*t* from a secondary bead (resp. *P*_*b*,4_ = λ_3_Δ*l*_3_Δ*t* from a tertiary bead), where Δ*l*_2/3_ is the bead interspacing for secondary (resp. tertiary) branches and λ_2/3_ is the branching rate of tertiary (resp. quaternary) branches. Whenever a new branch (called child) is created, we place the first bead of the newly created child branch are the coordinate location

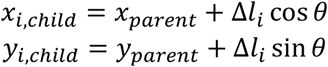

where (*x_parent_*, *y_parent_*) is the coordinate of the parent bead, the index *i* runs for the order of complexity of the branch and *θ* is chosen according to a uniform distribution on [0,2*π*].

#### Polymerization/depolymerization

We assume that branches are either in a polymerizing or a depolymerizing state. A branch in a polymerizing state grows with a rate *ν_i,on_*, where the index *i* again runs for the order of complexity of the extension. In this way, at each time step Δ*t* the number of beads of the branch is incremented by one. The position of the new bead *j* is:

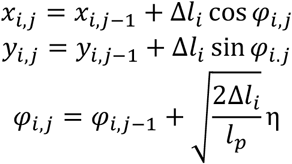

where (*x*_*i,j*−1_, *y*_*i,j*−1_) is the position of the previous bead in the branch, η is a centered unit Gaussian variable and *l_p_* is the persistence length of the dendrite branch. We considered a persistence length of *l_p_* = 17μ m, which is consistent with values measured in the actin bundles ^73^ (see Table 1).

Conversely, a branch in a depolymerizing state shrinks with a rate *ν_i,off_*. Experimental branch tracking suggest that we can set *ν_i,on_* = *ν_i,off_*. Depolymerization then occurs through removal of the tip branch bead. We prevent branch depolymerization beyond a forking bead. If a child was created from this bead, the parent branch changes to a polymerizing state and starts to grow again. A branch disappear if its last bead is removed through depolymerization.

#### Spontaneous switching

We define the rates *k_on_* (resp. *k_off_*) for branches to switch from a depolymerizing (resp. polymerizing) state to a polymerizing (resp. depolymerizing) one. Practically, in our simulation, we evaluate at each time step Δ*t* whether a branch should switch from a depolymerizing state to a polymerizing state; the switch occurs stochastically everytime we find that a random variable uniformly distributed on (0,1), denoted U, satisfies the relation:

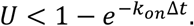

Therefore, the mean duration of polymerization phases read *τ_on_* = 1/*k_on_* .Similarly, the statistics of the switch from polymerization to depolymerization is evaluated according to a rate denoted *k_off_* (and *τ_off_*).

#### Contact-induced depolymerization

Whenever the distance between the last new bead of a polymerizing branch and any bead of other branches is smaller than a certain threshold *d_th_*, depolymerization is implemented with probability *p_off_*. The case *p_off_*= 1 corresponds to the control case; we find that *p_off_* = 0.3 provides a satisfactory fit of the *dscam^p1^* mutant case.

#### Aging

We implement a progressive reduction in the values of the dynamical parameters at regular time interval (corresponding to 2h experimentally). In particular, both the branching and switching rates, *λ_i_* and *k*_*i,on*/*off*_ respectively, are reduced by a factor *f_i_*, with *i* the order of complexity of the branch, while the polymerizing and depolymerizing rates, *ν*_*i,on*/*off*_, are reduced by a different factor *f_ν,i_*. At each reduction step, we remesh each branch with a new interspacing given by *ν_i,on_ f_ν,i_Δt* (with *ν_i,on_* the initial growth rate), which amounts to increasing the number of beads. The interplay between the possible values of *f_i_* and *f_ν,i_* defines the two aging protocols presented in the main text (see also Fig. 4d).

## Code availability

The custom codes used to process images analyze data and run simulations are available upon request.

---

**Figure S1.**
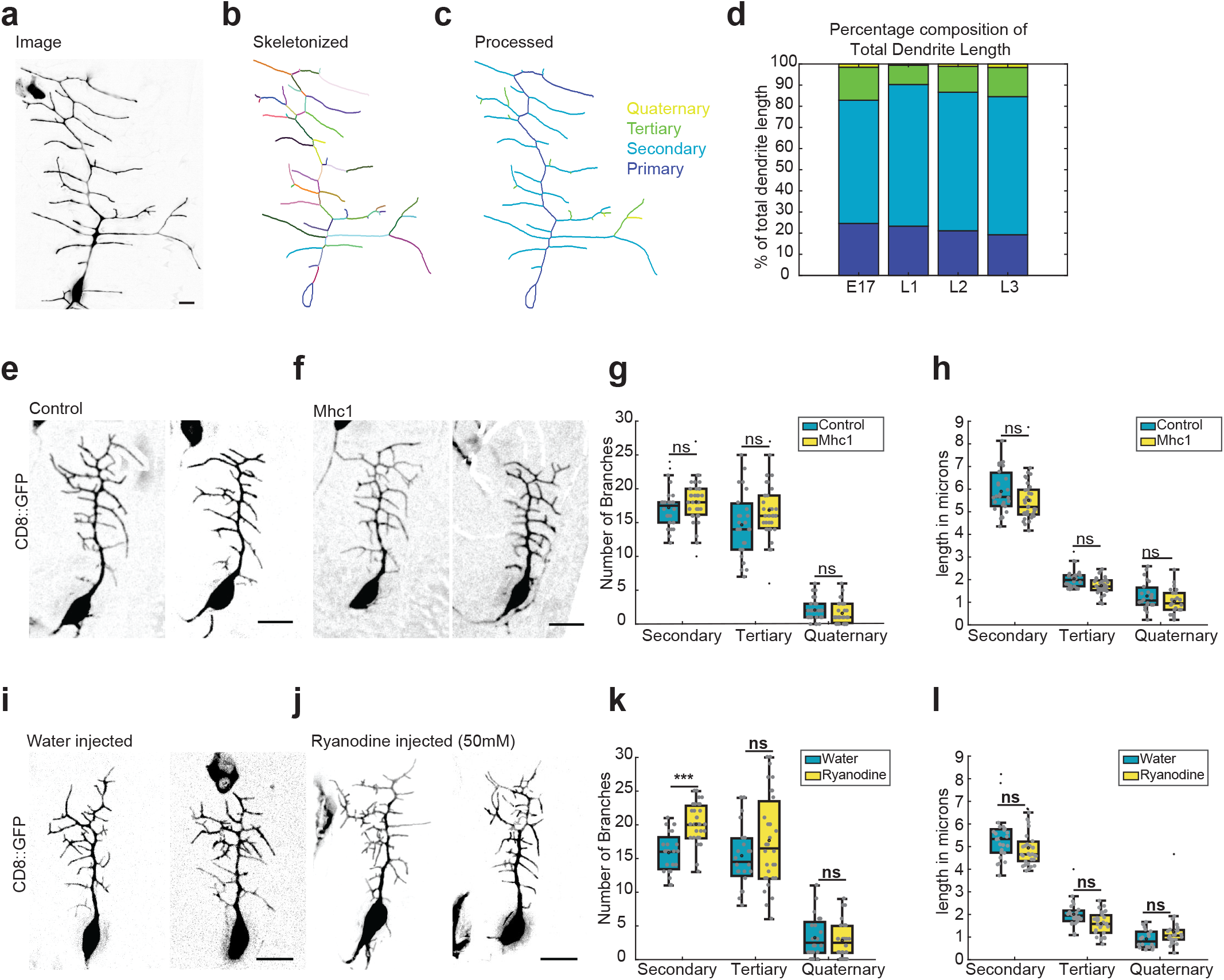
Branch ordering class I vpda neuron and blocking muscular contractions. (a) Class I vpda neuron at the 3^rd^ instar (L3 72h AEL±3) (b) Skeletonized neuron obtained from ‘Dendritic Arborization Tracer’. Every segment (length of dendrite between two vertices) has a unique identity represented by different colours. (c) Processed dendrite classified into 4 orders of complexity. Primary (Indigo), Secondary (blue), Tertiary (green), Quaternary (yellow). (d) Quantification of the percentage composition of total dendrite length. E17 n=26, L1 n=48, L2 n=50 L3 n=38 neurons (e) Control Class I vpda neurons taken from homozygous 2-21gal4/UAS-mCD8::GFP embryos (20h AEL±3). (f) Class I vpda neurons taken from 2-21gal4/UAS-mCD8::GFP, mhc1 mutant embryos (20h AEL±3). (g) Quantification of the number of branches at each order of complexity (h) Quantification of the branch lengths at each order of complexity. For (g)and (h) Control (Blue) n=27 Mhc1 mutant (yellow) n=35. (i) Control Class I vpda neurons injected with water (j) Class I vpda neurons injected with Ryanodine. For (i) and (j) embryos were taken from homozygous 2-21gal4/UAS-mCD8::GFP (k) Quantification of the number of branches at each order of complexity. (l) Quantification of the branch lengths at each order of complexity. For (k) and (l) Water (blue) n=24 Ryanodine (yellow) n=31. For each box, the central line is the median, the ‘o’ is the mean, the box extends vertically between the 25th and 75th percentiles, the whiskers extend to the most extreme data that are not considered outliers, and the outliers are plotted individually. Statistical significance has been calculated using Mann-Whitney U test. ns, * p<0.05; ** p<0.01, *** p<0.001. All the panels have the same orientation: dorsal at the top, anterior to the left. Scale bars = 10μm.

**Figure S2.**
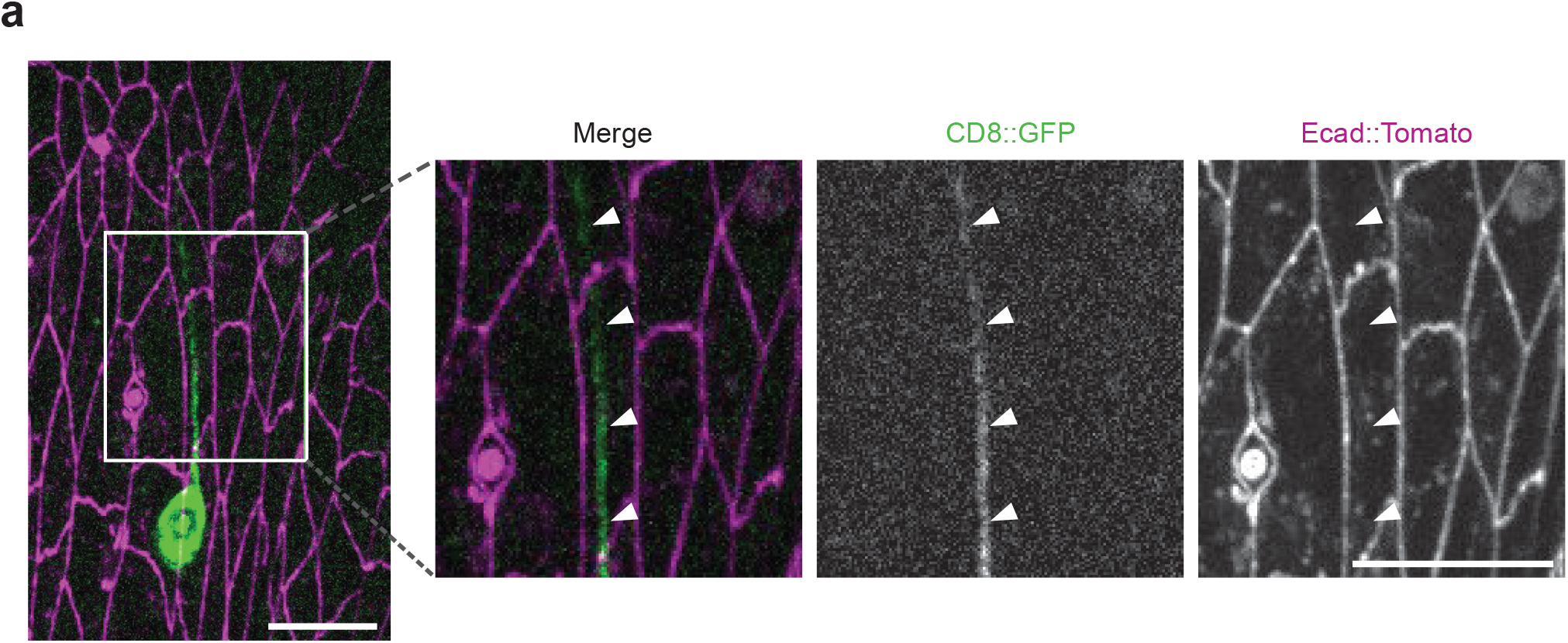
When the primary branch does not follow the stretched E-cadherin cell boundaries, it remains parallel to them. (a) Image of neuron at ~15h AEL co-imaged with Ecad::tomato. Inset: Close up of primary dendrite parallel to stretched E-cadherin cell boundaries as indicated by the white arrowheads. Scale bars = 10μm.

**Figure S3.**
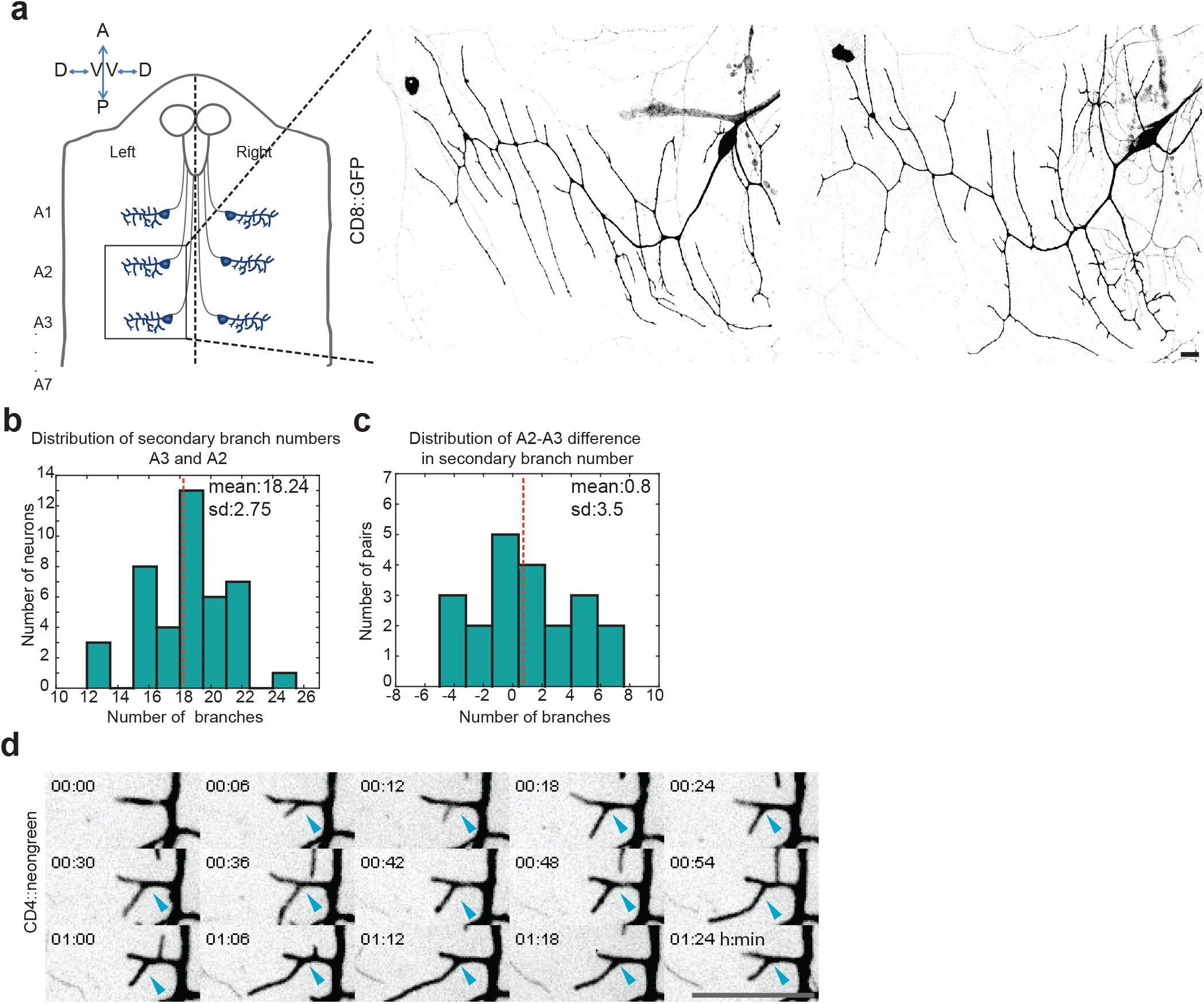
Difference between dendritic trees on abdominal segment A2 and A3. (a) Schematic: Larval filet (indicating vpda neurons in the abdominal hemi segments. Neurons in segment A2 and A3 on the left hemi-segment were used for analysis. Images of A2 (top panel) A3(bottom panel) vpda neurons at 96h AEL ±3 from homozygous 2-21gal4/UAS-mCD8::GFP larvae. The panel has the following orientation: Anterior at the top, Posterior at the bottom, ventral is the center and dorsal to the sides (b) Distribution of the total number of Secondary Branches n=42 (21 A2-A3 pairs). (c) Distribution of difference in number of Secondary branches between the A2-A3 pairs. (d) Secondary extensions do not shrink beyond forking points. Blue arrows indicate the forking point. (Movie7) Scale bars = 10μm.

**Figure S4.**
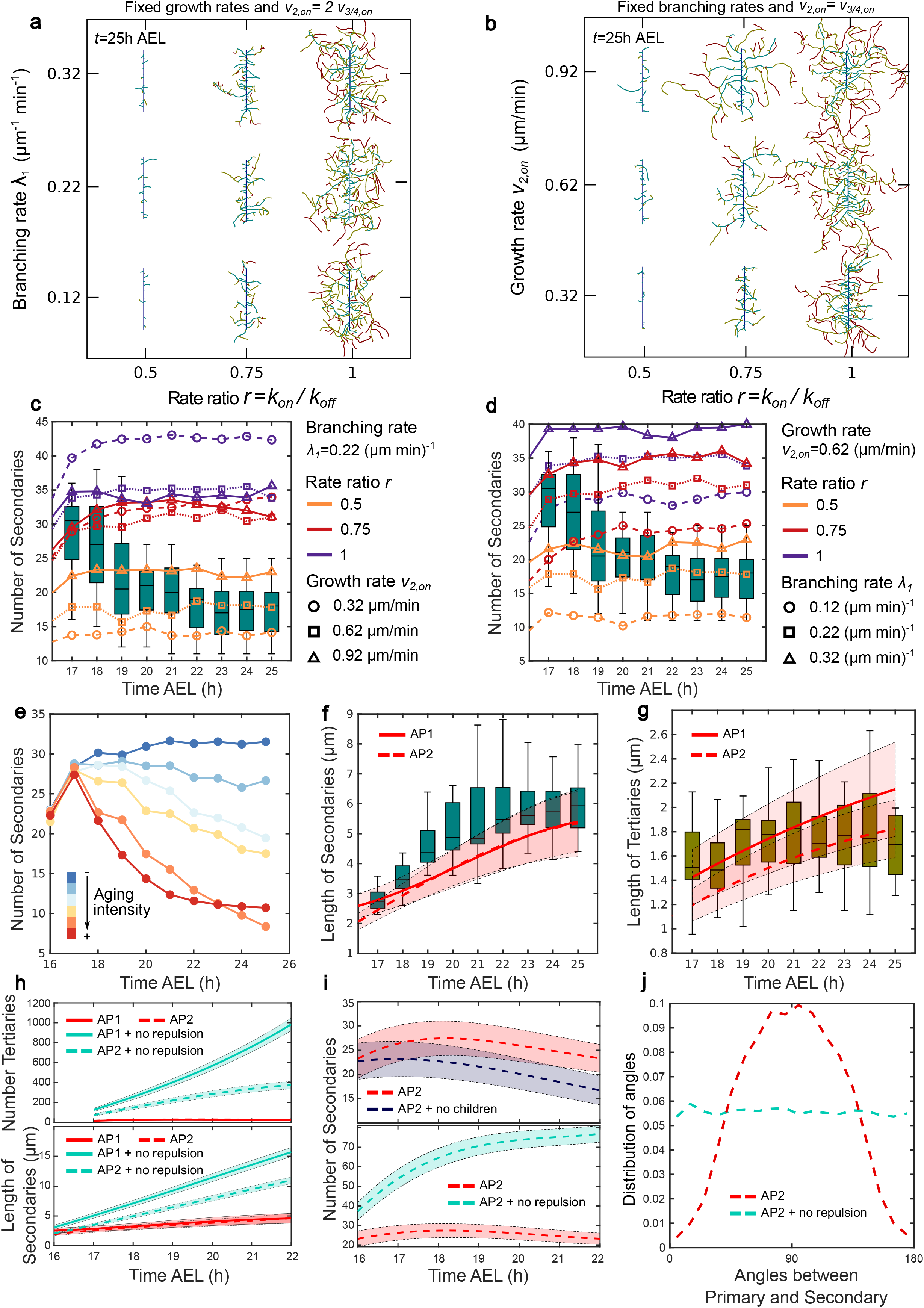
Computational Model for dendrite morphogenesis. Diagram of dendritic morphologies at 25h AEL for different values of the ratio of spontaneous switching rates *r* = *k_on_*/*k_off_* and branching rate of secondary branches *λ*_2,*on*_. The values of the other computational parameters are fixed and defined in Table 1, Option 1. Diagram of dendritic morphologies at 25h AEL for different values of the ratio of spontaneous switching rates *r* = *k_on_*/*k_off_* and growth rate of secondary branches *ν*_2,*on*_. The growth rates of tertiary and quaternary branches are given by *ν*_3/4,*on*_ = *ν*_3/4,*on*_ and the values of the other computational parameters are fixed and detailed in Table 1, Option 1. We again observe that the wide range of morphologies described by the model can be explored by varying the ratio *r*, as described in the Main Text for Fig. 4c. (c) and (d) Number of secondary branches over time from 16-25h AEL corresponding to the diagrams Fig. 4c (Main Text) and Fig S4a. In both figures, the growth rates for tertiary and quaternary branches are given by *ν*_3/4,*on*_ = *ν*_2,*on*_/2 and the other computational values are fixed and given by Table 1, Option 1. (e) Number of secondary branches over time from 16-25h AEL obtained in simulations with the same values of the parameters and different aging protocols. By increasing the aging intensity we can explore different dendrite developments which range from structures with a large number of branches to situations where only a few secondary branches survive at the end. (f) Average length of secondary branches and (g) Average length of tertiary branches over time from 16-25h AEL. Both in (f) and (g) red lines represents simulations with the two different aging protocols AP1 (full line) and AP2 (dashed line) (Fig. 4d Main Text) and the boxes represent the experimental data. (h) Top: Number of Tertiary branches over time from 17-22h AEL for the computational models with (red) and without (green) self-repulsion using both aging protocols AP1 (full line) and AP2 (dashed line) described in Fig. 4d of the Main Text. Bottom: Average length of secondary branches over time from 16-22h AEL for the computational models with (red) and without (green) self-repulsion using both aging protocols AP1 (full line) and AP2 (dashed line) described in Fig. 4d of the Main Text. (i) Top: Number of secondary branches over time from 16-22h AEL for the computational models with self-repulsion and child branches (red) and with self-avoidance but no child extensions (blue) using AP2 (Fig. 4d Main Text). Bottom: Number of secondary branches over time from 16-22h AEL for the computational models with (red) and without (green) self-repulsion using AP2 (Fig. 4d Main Text). (j) Probability distribution of the angles between primary and secondary branches at 22h AEL. Red and green lines represent simulations with and without self-repulsion, respectively using AP2 (Fig. 4d Main Text). The number of simulations for all cases in (c) and (d) and (e) was *N_sim_* = 50 while for all cases in (f), (g), (h), (i) and (j) it was *N_sim_* = 500. In all figures, simulation lines represent the average value of the realizations and shadows represent the 1*σ* confidence interval. In addition, a 2-point time-averaging was performed using spline interpolation for lines in (f), (g), (h) and (i). For (e), AP1 in (f) and (g), and AP1 with and without self-repulsion in (h) the initial values of the computational parameters are detailed in Table 1, Option 1. For AP2 in (f) and (g), AP2 with and without self-repulsion in (h), and all cases in (i) and (j) the initial values of the computational parameters are detailed in Table 1, Option 2. For each box corresponding to *in vivo* observations, the central line is the median, the box extends vertically between the 25th and 75th percentiles and the whiskers extend to the most extreme data that are not considered outliers.

**Figure S5.**
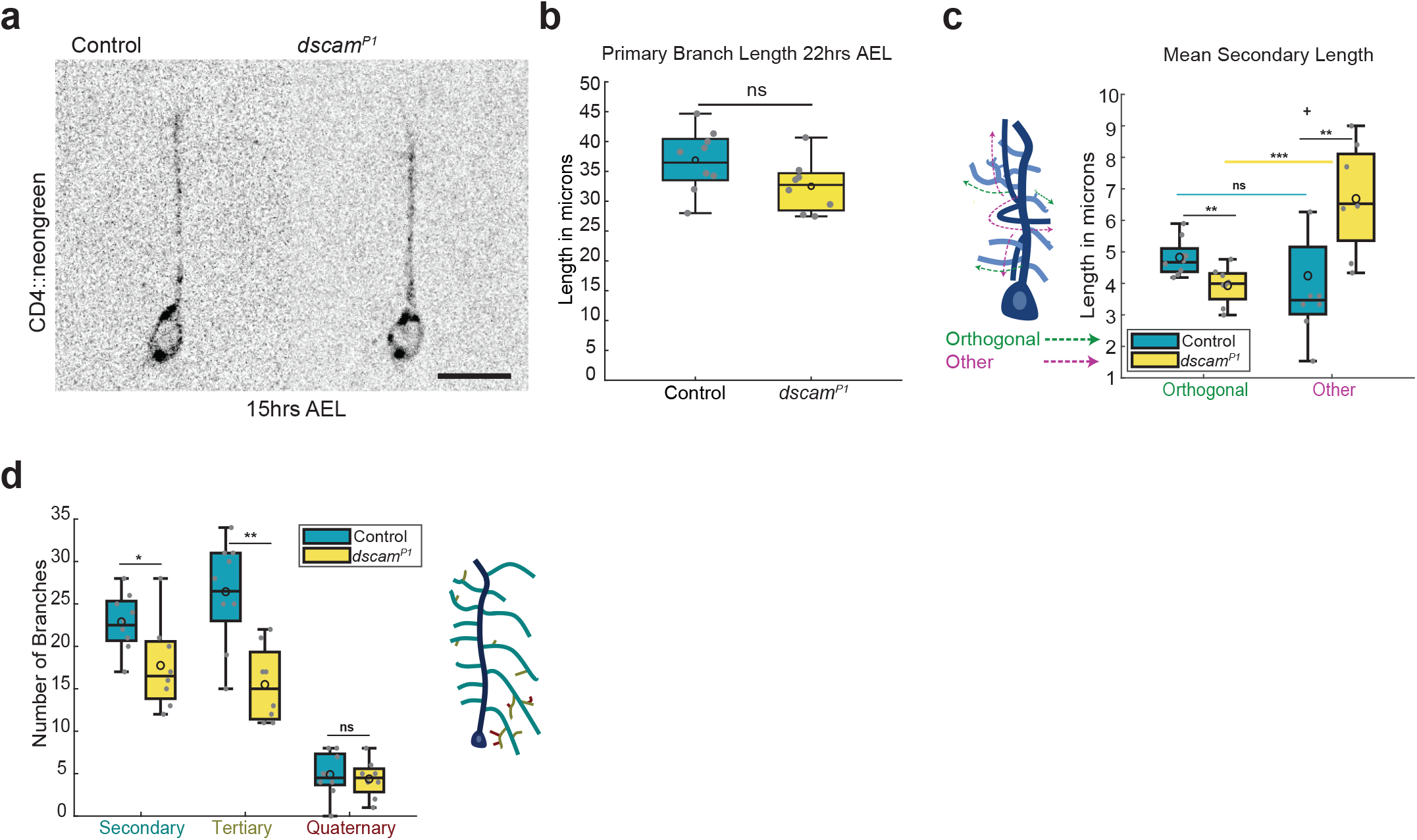
Secondary Branches require Dscam1 to allow a preferential orthogonal stabilization. (a) Left Panel: Control Class I vpda neurons taken from homozygous embryos of 2-21gal4/UAS-mCD4::neongreen. Right Panel: *dscam^P1^* mutant neurons taken from embryos of *dscam^P1^*/ΔDscam1;2-21gal4/UAS-mCD4::neongreen. Neurons imaged at ~15h AEL. (b) Quantification of Primary Branch lengths at ~22h AEL. (c) Schematic: Branches oriented outwards (Green arrows) Branches oriented in other directions (magenta arrows). Mean Lengths of Secondary branches oriented orthogonally vs non-orthogonally. (d) Quantification of number of branches at each order of complexity. Schematic: Primary branch (blue) Secondary branches (cyan) Tertiary (brown) Quaternary (red). Control (blue) n=9 *dscam^P1^* mutant (yellow) n=8. For each box, the central line is the median, the ‘o’ is the mean, the box extends vertically between the 25th and 75th percentiles, the whiskers extend to the most extreme data that are not considered outliers, and the outliers are plotted individually. Statistical significance has been calculated using Mann-Whitney U test. ns, * p<0.05; ** p<0.01, *** p<0.001. All the panels have the same orientation: dorsal at the top, anterior to the left. Scale bars = 10μm.

## Movie legends

**Movie1:** Time-lapse of embryonic development of the class I vpda neuron from ~16h to 25hrs AEL marked mCD8::GFP. Frame rate: 3min. Scale bars = 10μm

**Movie2:** Time-lapse primary dendrite development from ~14h to 15h AEL marked mCD8::GFP. Frame rate: 3min. Scale bars = 10μm

**Movie3:** Time-lapse of embryonic development of unablated vpda (control) neuron from ~16h to 25hrs AEL marked mCD8::GFP. Frame rate: 30min. Scale bars = 10μm

**Movie4:** Time-lapse of embryonic development of vpda neuron ablated at ~16h AEL marked mCD8::GFP. Frame rate: 30min. Scale bars = 10μm

**Movie5:** Time-lapse of vpda neuron ~18h AEL marked mCD8::GFP. Frame rate: 30sec. Scale bars = 10μm

**Movie6:** Simulation movie for two realizations for the parameter set defined in Table 1, Option 1 and aging protocol AP1. Primary branch (dark blue) has a constant length *L*_1_ = 30μm.

**Movie7:** Time-lapse of vpda neuron marked mCD8::GFP showing secondary dendrites do not retract beyond a forking point. Frame rate: 3min. Scale bars = 10μm

**Movie8:** Time-lapse of control vpda neuron development from ~17h-22h AEL marked mCD4::neongreen. Frame rate: 2min. Scale bars = 10μm

**Movie9:** Time-lapse of *dscam^P1^* vpda neuron development from 17h-22h AEL marked mCD4::neongreen. Frame rate: 2min. Scale bars = 10μm

**Movie10:** Time-lapse of *dscam^P1^* vpda neuron marked mCD4::neongreen showing remnant self-repulsion. Frame rate: 3min. Scale bars = 10μm

## Notes

### Competing Interest Statement

The authors have declared no competing interest.

### Summary of Updates

Supplemental Video files updated

